# Crowding drives terminal investment in a generalist pest

**DOI:** 10.64898/2025.12.12.694078

**Authors:** Shubha Govindarajan, Draupadi Patra, Deepa Agashe

## Abstract

Apart from genotype and environment, several fitness related traits of individuals are also influenced by the conditions experienced by their parents, i.e., by parental effects. To uncover the underlying physiological or molecular mechanisms, most studies analyze specific drivers in isolation, such as parental age or the degree of crowding experienced by the parents. However, potential interaction effects between different aspects of parental context are poorly understood. We analyzed the combined effect of parental age and population density on the number, fitness and physiology of offspring, in the red flour beetle *Tribolium castaneum*. Both factors independently reduce fitness, leading us to predict that old parents from high density should produce fewer offspring of the lowest quality. Surprisingly, older parents exposed to crowding produced more eggs than expected. These eggs were larger, had higher survival in an optimal as well as a suboptimal resource, and were provisioned with more lipids. This result is consistent with terminal reproductive investment in response to high density. The interaction between two ecologically relevant contexts — ageing and crowding — thus drives terminal investment via nutrient provisioning, perhaps contributing to the success of this generalist pest.

## INTRODUCTION

Offspring inherit genetic material as well as non-genetic provisions from their parents. This may vary depending on the context in which the parents developed or reproduced, leading to trans-generational parental effects (reviewed in (Edwards et al., 2021; Mousseau & Dingle, 1991; Räsänen & Kruuk, 2007; Roach & Wulff, 1987; Walsh et al., 2024; Wolf & Wade, 2009)). For instance, parental ageing typically reduces offspring number and quality (reviewed in Monaghan et al., 2020; Promislow et al., 2022), as in the parasitic wasp *Eupelmus vuilleti* where older mothers produced smaller eggs, resulting in offspring with low feeding rates and lower survival (Giron & Casas, 2003). However, such effects of parental age are not always monotonically negative. For instance, in case of terminal investment, adults increase their reproductive effort towards the end of their reproductive life, producing either higher quality or more offspring (Duffield et al., 2017; M. N. Muller et al., 2024). Apart from intrinsic factors such as ageing, the environment experienced by the parents — including competition, predation, nutrient availability, and environmental stress — also influences aspects of offspring fitness such as their size, development, survival and longevity (reviewed in (Burton & Metcalfe, 2014; Mousseau & Dingle, 1991; Rossiter, 1991, 2003)). For instance, crowded female damselfish *Pomacentrus amboinensis* produce smaller offspring (McCormick, 2006). Such parental effects are reported across many insects, plants, fish, birds, reptiles, and rodents (reviewed in Mousseau & Dingle, 1991; Rossiter, 2003). Thus, a large body of work shows pervasive and strong effects of parental context on offspring traits and fitness.

Broadly, two types of mechanisms may drive such parental effects. One class of mechanisms is context-dependent provisioning of nutrients, hormones, or immunological factors, which can directly or indirectly alter subsequent nutrient acquisition and metabolism in the offspring (reviewed in (Bebbington & Groothuis, 2021; Bonduriansky & Day, 2009)). For example, in the wasp *E. vuilleti* described above, offspring of older mothers have lower fitness because they receive less protein, sugar and lipid from the mother (Giron & Casas, 2003; D. Muller et al., 2017). In the case of the damselfish, crowding increases the levels of ovarian cortisol, which in turn reduces vitellogenin provisioning in eggs and causes offspring to be smaller (McCormick, 2006). Another class of mechanisms involves environment or experience-induced epigenetic changes such as DNA methylation (Becker & Weigel, 2012; Jablonka & Raz, 2009; Lim & Brunet, 2013) that are inherited by the offspring, altering gene expression patterns with downstream effects on offspring behaviour and fitness (Heard & Martienssen, 2014; Ruebel & Latham, 2020). For instance, exposure to sublethal doses of insecticide in the Colorado potato beetle (*Leptinotarsa decemlineata*) reduced global DNA methylation in the parental and F2 generations, and increased DNA methylation in the cytochrome P450 gene, which is involved in detoxification, thereby leading to insecticide resistance (Brevik et al., 2021). Although it is often difficult to identify the mechanisms underlying parental effects, this information is important to predict the magnitude and timescales on which they persist, and therefore, the potential evolutionary consequences of these effects (Marks & Lailvaux, 2024; Mousseau et al., 2009; Mousseau & Dingle, 1991).

Most studies typically focus on a single parameter such as age or competition, but in reality, several factors may jointly or independently generate parallel or opposing parental effects (Kim & Morales, 2025). For instance, we previously found that in the red flour beetle *Tribolium castaneum*, parental age and crowding interact to influence female oviposition behaviours and offspring traits in an optimal resource (wheat flour) as well as a suboptimal one (finger millet flour) (Ravi Kumar et al., 2022). Young females that experienced low density laid many more eggs, but their offspring developed more slowly compared to the offspring of young females from high density. Thus, females appear to optimize offspring number (i.e., offspring quantity) at low density due to low conspecific competition, but maximize offspring development rate (i.e., offspring quality) at high density, thereby optimizing fitness traits according to the context (Ravi Kumar et al., 2022). These results suggested that assuming a constant environment, females differentially regulate offspring traits to suit the likely context in which the offspring would develop. However, the physiological mechanisms underlying these effects remain unclear. This information is important because the net outcome for offspring fitness may depend on whether the interacting parameters alter offspring traits via similar or distinct processes.

Here, we addressed this question using a full factorial experiment with young vs. old red flour beetles that experienced low vs. high population density. We tested for their investment in offspring quantity (number of eggs) and egg quality (the size and nutrient composition of eggs, as well as egg survival and development rate in an optimal and a suboptimal resource). We find that females indeed optimize offspring performance in contexts aligned with their own experience, as observed in prior work (Kumar et al., 2022). Surprisingly, despite bearing the double cost of old age as well as crowding, old females from high density populations produced the best-quality eggs in terms of survival, likely via increased lipid provisioning. We suggest that this represents a case where two parameters interact to give rise to density-dependent terminal investment via nutrient provisioning. These parameters — ageing and competition — are ubiquitous aspects of life history, and may thus be broadly useful to understand parental effects in many species.

## METHODS

### Obtaining parents of different age and density contexts

We used an outbred stock population (described in Ravi Kumar et al., 2018)) consisting of ∼2000 adults maintained in wheat flour to obtain individuals of different age and density contexts for our experiments (Figure 1A). Three of the four parental contexts used in this study were generated as described earlier (Ravi Kumar et al., 2022), and the oldLD context is described here for the first time. The detailed protocol is shown in Figure S1. In brief, to obtain LD (low density) beetles, we isolated female pupae from large stock populations for 2 weeks post eclosion, allowed them to mate with single males of the same age for 4 days, and then removed the males. Then we used these females for our experiments, either right away (youngLD) or after allowing them to age until 6 weeks post-eclosion. We obtained HD (high density) beetles directly from a stock population maintained with 3 days of egg laying each generation to minimize age differences between individuals. We sampled beetles from this population at the appropriate age (2 weeks or 6 weeks post-eclosion), assuming that all individuals had already mated (Figure S1). We note that HD individuals likely vary in age (1-3 days), but this variation is much smaller compared to the difference between young and old age treatments (14 days vs. 28 days post eclosion).

**Figure 1:**
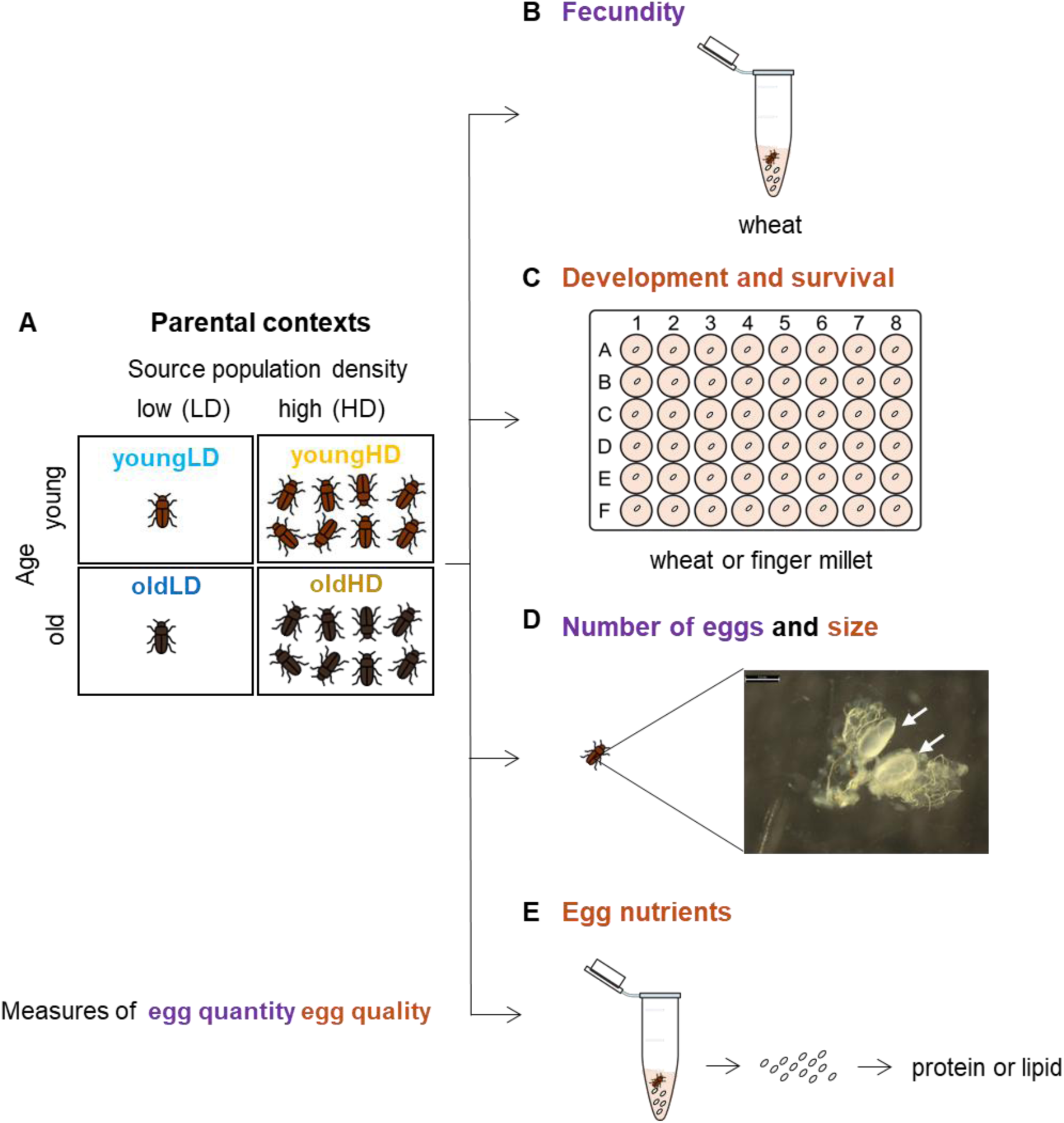
Schematics of the experiment design. **(A)**. The four parental contexts used in our experiments: young low density (youngLD), young high density (youngHD), old low density (oldLD), and old high density (oldHD). Mating occurred between adults from the same context. Fig S1 shows further details. **(B–E)** Schematics showing assays to quantify parental effects on offspring quantity (purple) and quality (orange). **(B)** Fecundity **(C)** Survival and development rate of eggs reared in isolation **(D)** Number and size of eggs in ovaries of dissected females. The image is a sample photograph with arrows pointing to eggs. Scale bar = 0.5 mm. **(E)** Nutrients deposited by females in clutches of eggs.

### Measuring egg quantity: Fecundity assays

We allowed mated females from each context to oviposit individually in 1 g of double sifted wheat flour in a 1.5 ml microcentrifuge tube for 48 hours, and counted the eggs laid (n = 50 females, Figure 1B). Sifting the flour twice removes large flour particles that interfere with egg collection. We froze the females at -80°C for further analysis, and used the collected eggs for egg quality measurements (see below).

### Measuring egg quality: Survival and development rate

We measured egg quality in both wheat and finger millet, with eggs developing individually in the absence of intraspecific competition and cannibalism. For this assay, we pooled the eggs laid by all females from a specific context (see above), and distributed 96 randomly chosen eggs in wells of 48-well microplates (Costar) containing ∼0.6 g of resource (wheat or finger millet) (Fig 1C). We covered plates with Parafilm, made a small hole above each well for ventilation, and incubated at 33° C (Figure 1C). After 3 weeks, we noted the total number of live individuals (measure of survival), and the proportion of eggs that had successfully pupated (measure of development rate). A higher proportion of pupae indicates faster development.

### Measuring egg quality: Number and size of eggs

From the frozen females from fecundity assays (see above), we dissected 30 females per context under a stereomicroscope (Leica Application Suite v 4.1.2) at 4x ocular lens magnification, to measure the number and size of eggs present in the ovaries. We thawed each female, rinsed in 70% ethanol followed by a rinse in autoclaved saline (0.9% NaCl), and placed it on a 5% agar plate and cut the tip of the abdomen using fine scissors (Fine Science Tools). The ethanol rinse was required only because we used the dissected eggs for another project where we analysed the egg microbiome (Sanjenbam et al., 2025). We applied pressure to the sternum to force out the abdominal contents. We teased out the ovary on a glass slide with a drop of saline and counted the number of mature eggs. Using the Leica application, we imaged each female’s body contents (Figure 1D). To measure egg size, we used the “Object Area” feature to quantify the total area of each egg from the images, and calculated the average egg size for each female.

### Measuring egg quality: Nutrient provisioning

For this experiment, we generated a new set of adults from each of the four contexts. We followed the fecundity assay protocol described above, counted and transferred all eggs from a given female to a fresh tube, and stored them at -80°C until further analyses (n=12 females per context per nutrient, i.e., egg clutches from distinct subsets of females from a given context were used to measure each nutrient, Figure 1E).

To estimate protein content, we homogenized all eggs from a clutch with a micropestle in 200 µL phosphate-buffered saline (PBS) containing 3 µl protease inhibitor cocktail (Sigma-Aldrich, P8340). After centrifugation at 12400 rpm for 12 minutes, we quantified total protein in 25 µl of the supernatant, with a BCA assay kit (G-Biosciences Cat #786-570). We prepared a set of standards using bovine serum albumin (G-Biosciences) diluted in PBS (0, 0.2, 0.4, 0.6, 0.8, 1 µg/µl), and added 25 μl of each standard to 200 μl BCA reagent. We incubated samples and standards in a 96-well plate (Costar) at 37°C for 30 minutes, and measured the absorbance at 562 nm using a microplate reader (Biotek Epoch 2). We obtained a standard curve of the absorbance as a linear function of the concentration of the standards, and used it to determine protein concentration of each sample.

We measured total lipid content of each egg clutch using chloroform–methanol extraction followed by a phospho-vanillin colorimetric assay (Van Handel, 1985). We homogenized each sample in 200 µL of a 1:1 (v/v) chloroform–methanol mixture and collected 50 μl of the supernatant after centrifugation at 12400 rpm for 12 minutes. We prepared 50 µL lipid standards (0.2, 0.4, 0.6, 0.8 and 1 µg/µl soyabean oil in the chloroform methanol mixture). We evaporated the solvent by heating the tubes at 60 °C for 30 minutes and then added 20 µl of concentrated H₂SO₄ (98% w/w, 18 M). After 10 min of heating at 100 °C in a heating block, we cooled the tubes to room temperature. We added 480 µl of phospho-vanillin reagent (prepared by dissolving 600 mg of vanillin (Sigma Aldrich Cat #V1104-2G) in 100 ml water at 60°C and adding 400 ml of 85% orthophosphoric acid (Qualigens Cat # G29245)) and incubated tubes at 37 °C for 10 min in the dark to allow color to develop. We recorded absorbance at 530 nm using a microplate reader (Biotek Epoch 2), to quantify total lipid content using the standard curve.

We calculated total protein and lipid in each sample by multiplying the measured concentration by the total volume of solvent used for homogenization. We determined protein and lipid per egg by dividing the total values measured for the entire egg clutch per female, by the number of eggs in the clutch.

In a pilot experiment, we measured the trehalose content of 2 groups of 10 eggs from each of the four female contexts, by incubating homogenized samples with trehalase enzyme (Sigma Aldrich, T8778) to break down trehalose to glucose and galactose, and assaying the glucose formed using the glucose oxidase assay, as described previously (Gupta et al., 2019; Gupta & Laxman, 2020). However, we did not detect any glucose (and hence trehalose) in any of the samples, whereas the standards (0, 20, 40, 60, 80 and 100 µg/ml of glucose in water) gave the expected values of glucose. Hence, we did not conduct any further measurements of trehalose in this study.

### Measuring context-specific nutrient content of adults

To test whether oldHD adults had distinct body condition from youngHD adults (potentially explaining their greater investment in eggs), we measured the protein and lipid content of adults from youngHD and oldHD contexts using the same procedure described for eggs. Individual adults were ground up with liquid nitrogen and suspended in 500 µl of PBS (for protein) or chloroform-methanol extraction solvent (for lipid).

### Data analysis

All data were analyzed in R v4.4.3 (R Core Team, 2025). We used a linear model to analyze the effect of female age and density on each of the response variables measured (fecundity, number of eggs in the ovary, average egg size, and protein and lipid at the clutch and single egg level). We used a generalized linear model (GLM) with binomially distributed errors to analyze the effect of female context and resource (wheat or finger millet) on mortality and development rate. To compare mortality and development between two resources for a given female context, we used the chi-square test, and applied a Hochberg correction to account for multiple hypothesis testing. In cases where the success (or failure rate) was very low, we used a Fisher’s exact test. We applied an ANCOVA (Analysis of covariance) to test whether protein and lipid investment per egg was correlated with fecundity, and whether this relationship differed among the four contexts or as a function of parental age and density.

We used the packages “ggplot” (Wickham, 2016), “ggpubr” (Kassambara, 2023) and “gghalves” (Tiedemann, 2022) for data visualization, “dplyr” (Wickham et al., 2023) and “tidyr” (Wickham et al., 2024) for summary statistics, and “lmerTest” (Kuznetsova et al., 2017), “emmeans” (Lenth, 2025), “effectsize” (Ben-Shachar et al., 2020), “car” (Fox & Weisberg, 2019), EnvStats (Millard, 2013) and “multcompView” for data analysis (Graves et al., 2024).

## RESULTS

### Ageing and high density reduce fitness, but oldHD parents perform better than expected

Prior work lacking a full-factorial design showed that females from different age or density contexts had different fecundity (Ravi Kumar et al). Our primary goal in this work was to test for interaction effects and determine the physiological basis of these differences. As expected, young females that experienced low density (youngLD females) had the highest fecundity, laying 33 eggs on average in 48 hours (Figure 2A). Supporting previous observations, high density reduced fecundity in females of both age groups (Figure 2A, Table S1). We also found interaction effects: ageing reduced fecundity, but only for parents that experienced low density (Figure 2A; ANOVA *p_age_*<0.001, *p_density_*<0.001, *p_ageXdensity_*<0.001, Table S2). Thus, despite experiencing both ageing and higher density, oldHD females had similar fecundity as youngHD females (average of 10 and 12 eggs respectively), suggesting that oldHD females somehow mitigated these doubly detrimental effects.

**Figure 2:**
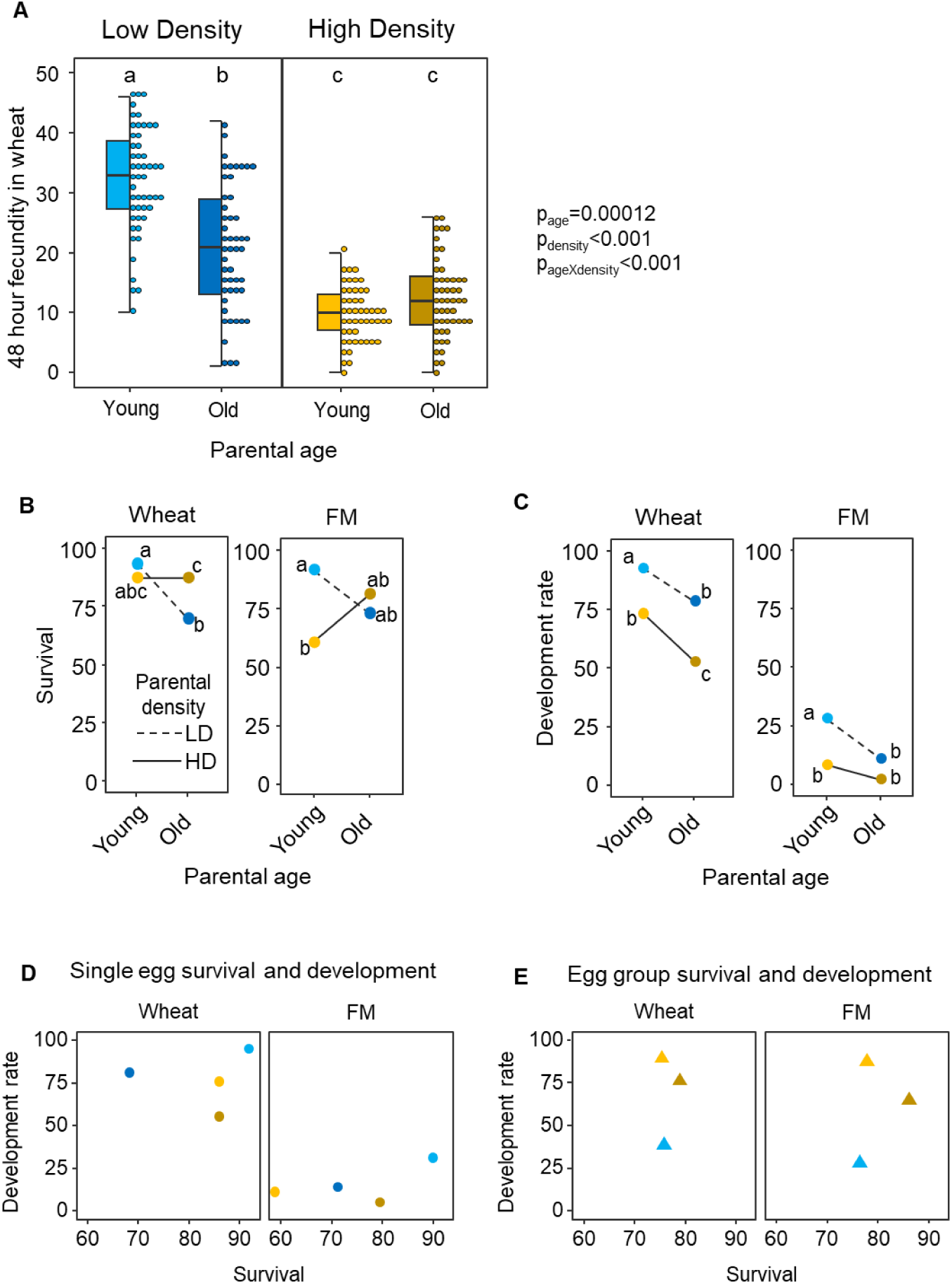
Parental age and experienced density affect fecundity and egg quality. (**A**) Number of eggs laid by females from each of the four contexts in wheat flour in 48 hours. Boxplots represent the median and interquartile range. Each point shows fecundity of a single female. (**B–C**) Measures of egg quality over 3 weeks of development: percentage of eggs that survived, and proportion of surviving offspring that pupated successfully (development rate) as a function of parental age, for offspring reared in (**B**) Wheat or (**C**) Finger millet (FM). Females only laid eggs in wheat, and we placed them in the appropriate resource. Each point represents the mean trait value, and is coloured as indicated in panel A. (**D–E**) Development rate vs. survival in each resource for (**D**) Eggs that developed singly (replotted from panels B and C) and (**E**) Eggs that developed in sibling groups (reproduced here from the wheat-only control of Experiment 2 in Ravi Kumar et al., 2022). In panels A–C, letters indicate the results of pairwise comparisons, such that contexts sharing the same letter are not significantly different from each other.

Next, we tested whether egg quality differed across the four contexts, focusing on egg survival and rate of development. We collected eggs from the above assay and allowed them to develop individually, in the absence of conspecific competition or cannibalism. For offspring of LD parents, egg survival in wheat decreased with parental age, but did not change with ageing for HD parents (Figure 2B, Table S3, Table S4), mirroring the observed pattern for fecundity. Results for eggs placed in finger millet were also consistent with a mitigating effect in oldHD parents, whose offspring had survival similar to youngHD parents (Figure 2B, Table S4). Parental exposure to high density reduced offspring development rate in wheat, but parental age had no effect (Figure 2C, Table S3, Table S4). In finger millet, offspring development rate was higher in the offspring of youngLD parents, but did not differ across the other parental contexts (Figure 2C, Table S4).

Visualizing these two axes of egg quality together (survival and development rate), we observed a consistent pattern of higher-than-expected quality of offspring produced by oldHD females (Figure 2D). Overall, as anticipated, youngLD females laid the most and best quality eggs with the fastest development and highest survival in both wheat and finger millet (Figure 2D). With ageing, their eggs paid a large survival cost and a small development rate cost (compare light vs. dark points in Figure 2D). Exposure to high density led to relatively small costs along both axes in wheat, and a large survival cost in finger millet (compare light blue and yellow points in Figure 2D). Given these independent costs of ageing and density, we had anticipated that offspring of oldHD parents should have the lowest quality and pay the highest costs. Surprisingly, eggs of oldHD parents paid a survival cost that was either similar to youngHD (in wheat) or much lower (in finger millet), with a comparable or slightly higher cost of development rate (compare yellow and brown points in Figure 2D). Therefore, oldHD parents generally appeared to have better-quality eggs than expected given that they experienced both ageing as well as high density.

A prior study had also measured these outcomes for eggs from three of the parental contexts analyzed here; but those eggs were reared in sibling groups (Ravi Kumar et al., 2022; reproduced here in Figure 2E). Comparing results from both studies (Figure 2D vs. Figure 2E), we see that eggs of HD parents have faster development and/or lower mortality when competing with siblings; whereas eggs from LD females paid a higher cost when reared under competition. These results suggest that HD parents optimize offspring performance in HD, whereas LD parents optimize offspring performance in LD. Thus, parental context appears to determine which aspect of egg quality is maximized, under conditions consistent with parental experience. Notably, offspring of oldHD parents have overall comparable quality to those of youngLD parents, potentially trading off slower development rate with higher survival (Figure 2E; Ravi Kumar et al). Thus, oldHD parents consistently produce more and better quality offspring than expected, regardless of offspring rearing conditions (group size or resource).

### oldHD females produce larger eggs containing more lipids

To understand the physiological basis underlying the higher-than-expected quality of oldHD eggs, we dissected the ovaries of females from each context. All of them had a similar number of eggs (*p_age_*=0.5037, *p_density_*=0.3788, *p_ageXdensity_*=0.5886, Figure 3A, Table S5), implying that the observed fecundity differences (Figure 2A) arise due to differential rates of oviposition across female contexts. Eggs of oldHD females were larger (Figure 3B, Table S5), but the number and size of eggs were not correlated (Figure S2; Table S6; p_context_=0.0205, p_egg number_=0.2644, p_contextXegg number_=0.4737). Therefore, egg size is not altered commensurate with changing egg production, suggesting the lack of a simple tradeoff between egg number and quality (McIntyre & Gooding, 2000). Thus, oldHD eggs are larger even though they produce a similar number of eggs as youngHD females. Are the larger size and better quality a result of greater nutrient provisioning in these eggs? To address this question, we measured nutrients in eggs laid by females from different contexts.

**Figure 3:**
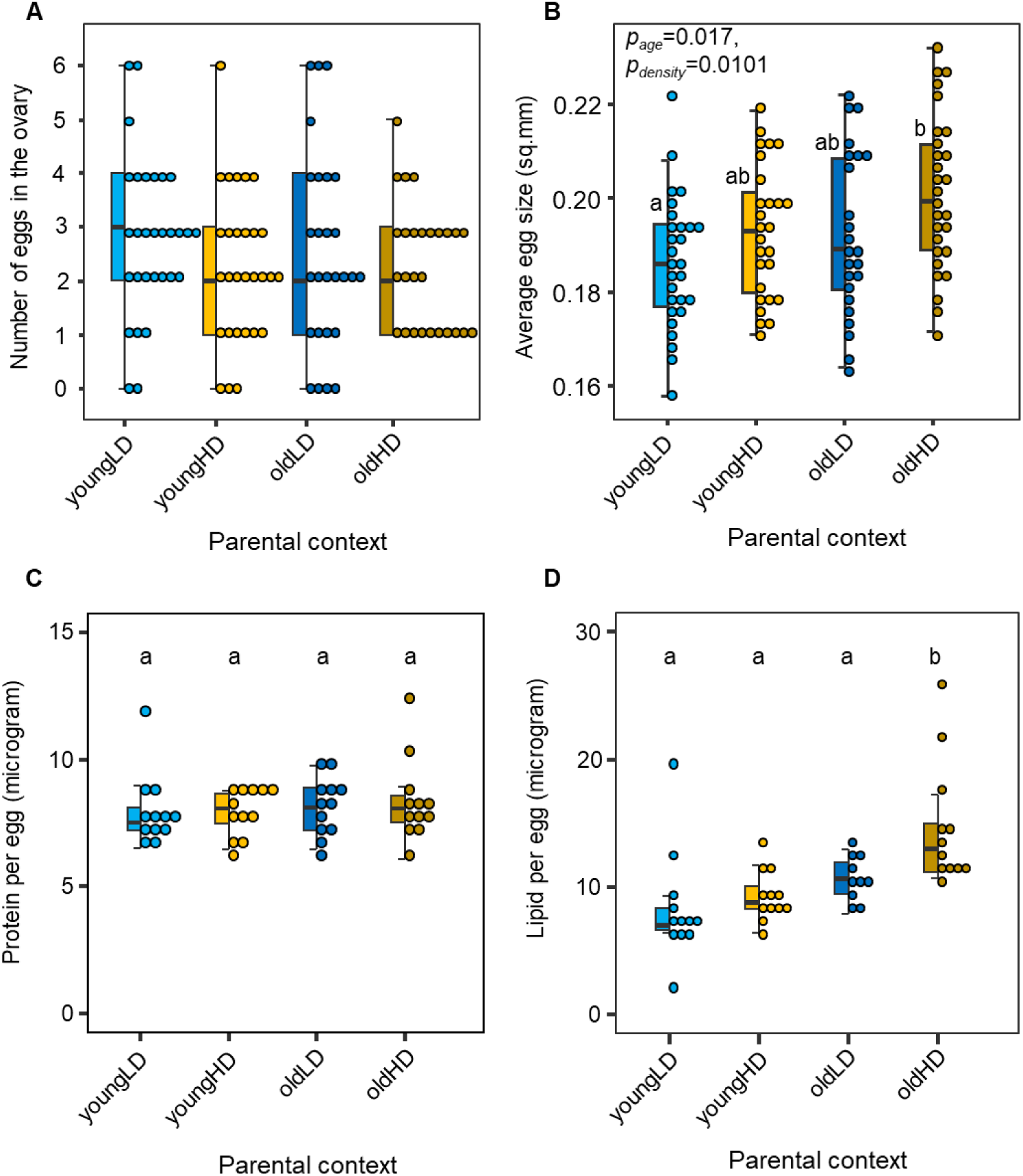
Number, size, and nutrient content of eggs from each parental context. (**A**) Number of eggs in the ovaries of dissected females from each context. Each point shows the number of eggs found in a single female. (**B**) Average size of eggs in the ovary of each female. (**C–D)** Boxplots (medians and interquartile range) showing (**C**) protein and (**D**) lipid per egg in the four female contexts, estimated from clutches of eggs. Letters denote the results of pairwise comparisons across contexts (ANOVA followed by post-hoc Tukey’s tests), such that contexts sharing the same letter are not significantly different from each other.

We could not detect any trehalose in eggs, so we focused on lipids and proteins as the major nutrient classes of interest. Females from all contexts invested similar amounts of total protein per clutch (Figure S3A-B), as well as per egg (Figure 3C). However, the amount of lipid investment at the clutch level and per egg varied by female context (Table S7 and S8), with a significant interaction between age and density at the clutch level (Table S7). Eggs of oldHD females had the highest lipid content, consistent with their overall higher quality and size (Figure 3D). Overall, both protein and lipid investment per egg declined with female fecundity (Figure S3C-D, Table S9): as females laid larger egg clutches, each individual egg received fewer provisions. However, for lipids, the slope of this relationship became steeper with age (Figure S3D). Thus, older females appear to be more constrained in how much they can provision their offspring. In contrast, the relationship between protein and fecundity was invariant (Figure S3C). Notably, despite differential lipid investment by youngHD vs. oldHD females in eggs, the females themselves had similar total body lipid content (Figure S4B, Table S10).

## DISCUSSION

Reproductive allocation is shaped by a fundamental life-history trade-off between investing in current offspring and preserving resources for future reproduction. Terminal investment theory predicts that when the expectation of future reproduction declines — through aging, declining body condition, or environmental stress — individuals should increase investment in current offspring to maximize remaining fitness (Clutton-Brock, 1984; Williams, 1966). Empirical work has typically detected this response through offspring quantity, such as increased egg number or fecundity late in life or under threat (Creighton et al., 2009; Duffield et al., 2017; Foo et al., 2023; Heinze & Schrempf, 2012; Jehan et al., 2021, 2022; Lemaître et al., 2015). However, recent studies show that terminal investment may also manifest through offspring quality, for example via increased egg protein content in response to a high pathogen load in *T. castaneum* (Schulz et al., 2023) or behavioral investment such as the number of offspring weaned in mammals (Lemaître et al., 2015).

We examined how parental age and population density jointly influence parental investment in *T. castaneum*. We asked whether aging parents adjust resource allocation between offspring number and quality depending on their environmental context, and found evidence for both. Consistent with terminal investment in offspring quality, older females exposed to high density produced larger eggs with more lipids, resulting in offspring with higher survival. At the same time, their fertility was similar to that of youngHD females, indicating terminal investment to minimize the cost of leaving fewer offspring. Importantly, this response was density dependent: under low density, offspring survival declined with maternal age, but egg provisioning did not change; whereas under high density, older females increased nutrient investment and offspring survival. These patterns suggest that younger females might be conservative in stressful environments — limiting reproduction when there is a possibility that future conditions might improve — but make a terminal investment later in life, when prospects for future reproduction diminish. As a result, females optimize offspring performance for the density that they experienced themselves, which may be advantageous when conditions are likely to remain constant across generations.

By integrating nutrient and demographic data, our results demonstrate that terminal investment can be multidimensional, with its expression modulated by environmental stress. *T. castaneum* females appear to integrate their internal state and environmental stress to dynamically adjust reproductive effort. By increasing lipid provisioning late in life under density stress, females enhance offspring survival. This density-modulated, nutrient-specific terminal investment shows how environmental stress can invert typical age-related declines in reproductive output, and mirrors context-sensitive patterns reported in other taxa. For instance, in the burying beetle *Nicrophorus orbicollis*, older females increased parental care at the expense of somatic maintenance when resources were scarce (Creighton et al., 2009). In *T. castaneum*, prior work found infection-dose-dependent trade-offs between maintenance and reproduction: severe infection elicited elevated egg protein content despite reduced fecundity (Schulz et al., 2023). Similarly, in *Daphnia magna*, greater maternal lipid provisioning by older females enhanced offspring mitochondrial function and longevity (Anderson et al., 2022), reinforcing the idea that maternal lipid investment may be a key axis of adaptive plasticity with far-reaching fitness implications. Our study thus underscores that insects can employ flexible biochemical allocation strategies to balance survival, reproduction, and offspring quality across changing environments.

In closing, we discuss some avenues for further research, and broader implications of our results. Our nutrient measurements were coarse-grained, but a detailed analysis of lipids and proteomes (such as that conducted in Prasad et al) may help understand exactly how lipid provisioning translates into higher offspring fitness. Another open question is whether our results reflect maternal, paternal, or combined effects. Future studies could independently manipulate maternal and paternal context to address this gap. Importantly, the broad patterns of egg quality differences across parent contexts were similar in both wheat and finger millet, suggesting that parental effects modulate intrinsic egg quality independent of available resources. We speculate that such generalized modulation of egg quality may help explain the flour beetle’s success as generalist pests of cereal flours. Another important implication is that such parental effects — via differential nutrient provisioning and resulting wide variation in offspring traits — could influence population dynamics (Benton et al., 2008; Plaistow & Benton, 2009), behaviour (Dias & Ressler, 2014), and life history traits (Plaistow et al., 2015). Therefore, to better predict population performance in a novel environment, it may be important to consider parental context and its underlying mechanisms (Catford et al., 2022).

## ACKNOWLEDGEMENTS

We thank Sunil Laxman and Sreesa Sreedharan for sharing their protocol and reagents for nutrient assays, Kanishk Tomar and Akshita Sohm for laboratory assistance, and Pratibha Sanjenbam, Adrita Chakraborty, Basabi Bagchi and other members of Adaptation Lab for feedback and comments on the manuscript. We acknowledge funding and support from the National Centre for Biological Sciences and the Department of Atomic Energy, Government of India (Project Identification No. RTI 4006 to DA and graduate student fellowship to SG).

## AUTHOR CONTRIBUTIONS

DA and SG conceptualized the study, SG and DP conducted experiments and analyzed data; DA acquired funding; all authors wrote the manuscript.

## Notes

### Competing Interest Statement

The authors have declared no competing interest.

