## Supplementary Information for "Crowding drives terminal investment in a generalist pest"

**SUPPLEMENTARY MATERIAL**: **Crowding drives terminal investment in a generalist pest**

**SUPPLEMENTARY FIGURES**

**Figure S1: Procedure to obtain adults of each of the four contexts for experiments**. For LD contexts, pupae were sexed and paired for mating within the same context. The pupae for LD contexts and adults for HD contexts were obtained from two different replicate stock populations.

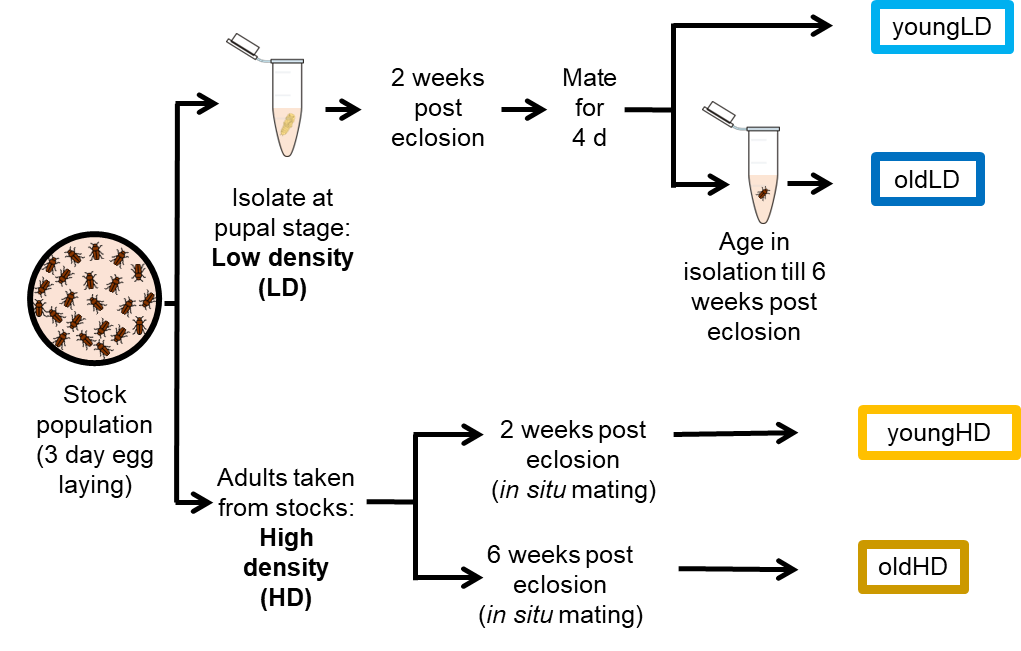

**Figure S2: Egg size is not correlated with number of eggs in the ovary.** Average egg size vs. number of eggs in the ovary of each parental. Error bars represent the standard deviation.

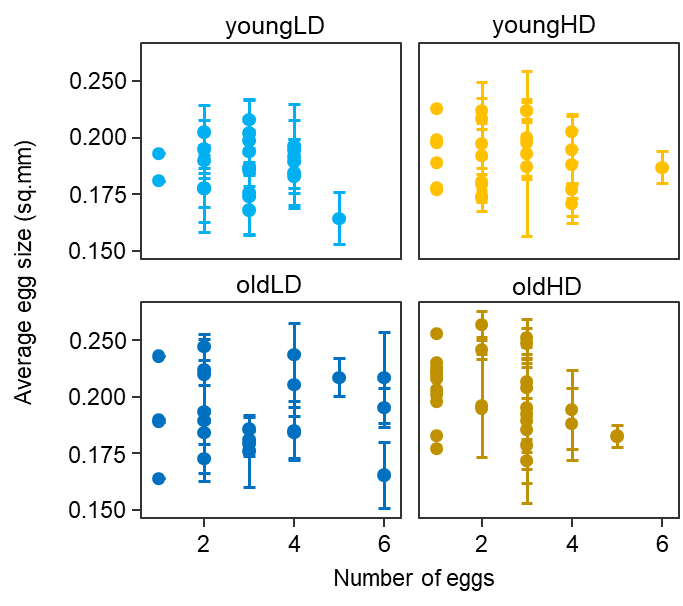

**Figure S3: Effect of parental context on clutch protein and lipid content**. (**A–B**) Boxplots show clutch level (**A**) protein and (**B**) lipid. Letters above the boxplots signify the results of a linear model with post-hoc Tukey corrections, such that boxplots sharing the same letter are not significantly different from each other. (**C–D**) Nutrients per egg, as a function of total parental fecundity (clutch size), for (**C**) protein and (**D**) lipid. Correlation coefficients displayed are Spearman’s rank correlations. In cases of significant correlations, linear regression lines fitted to the data are shown.

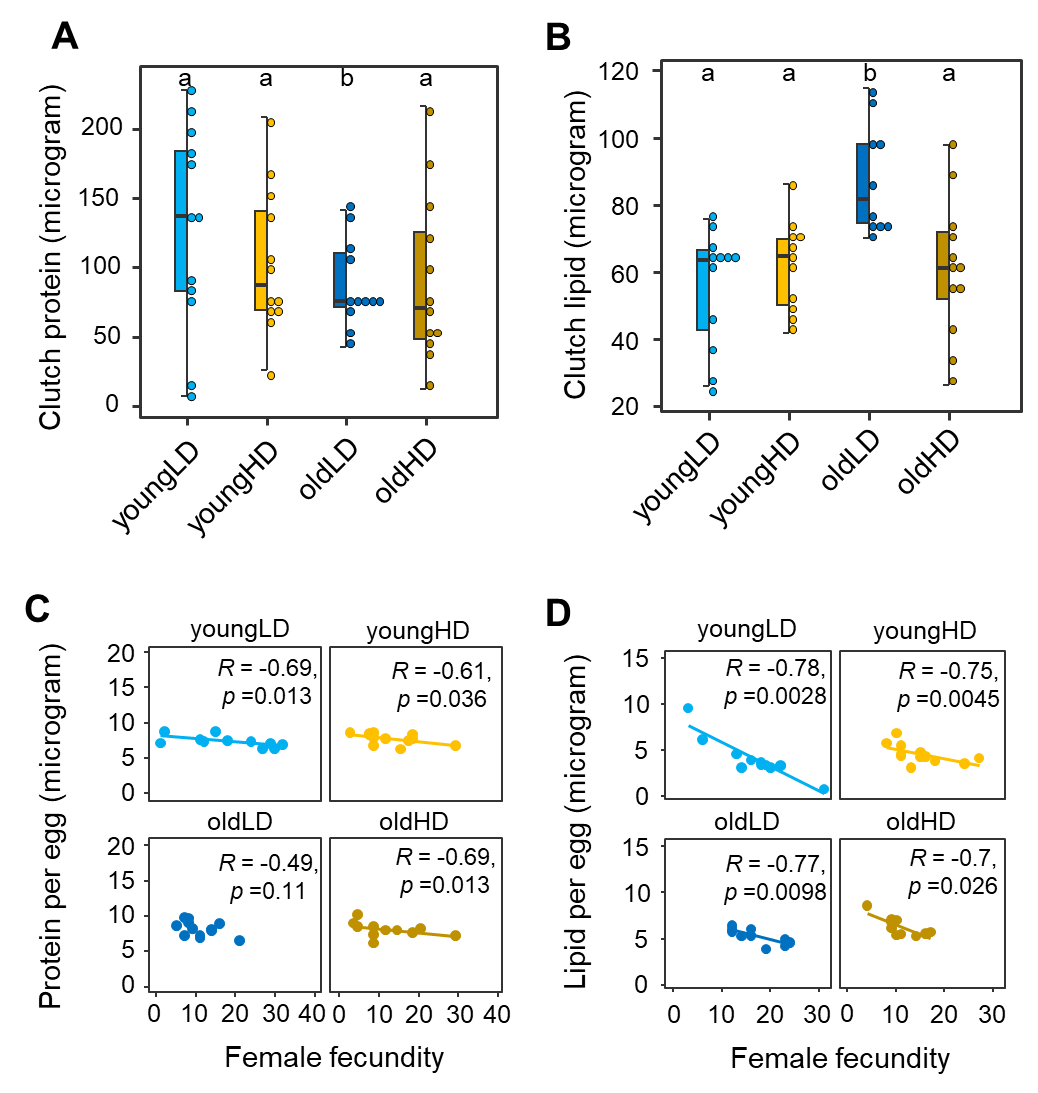
**Figure S4: Protein and lipid content are similar in young and old HD adults.** Boxplots of (**A**) protein and (**B**) lipid in youngHD (yellow) and oldHD (dark gold) adults. Letters above boxplots signify the results of a linear model with post-hoc Tukey corrections, such that boxplots sharing the same letter are not significantly different from each other.

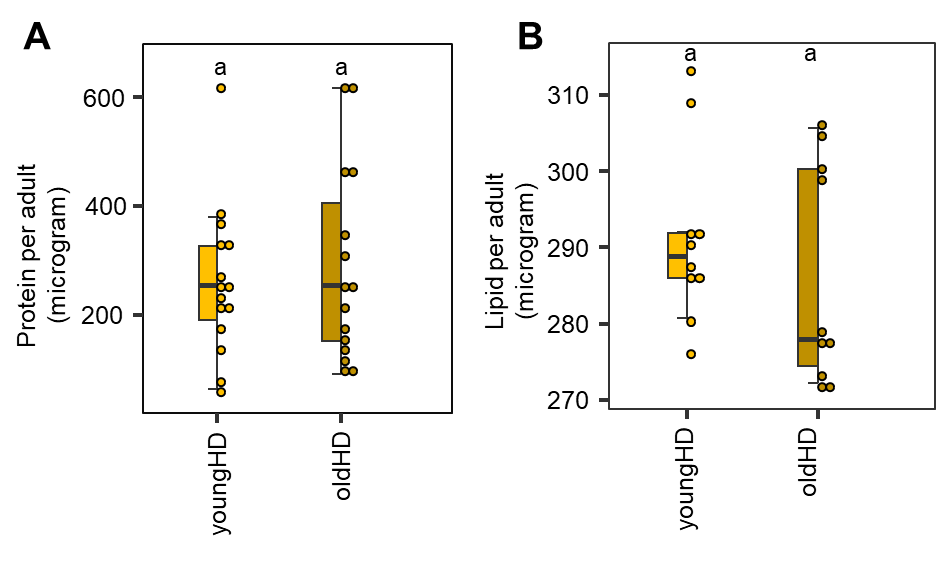

**SUPPLEMENTARY TABLES**

**Table S1**: Pairwise comparisons across contexts from the model shown in Table S1, with post-hoc Tukey’s correction. Significant values (p < 0.05) are highlighted in bold.

| Comparison | p-value |
| --- | --- |
| youngHD vs oldHD | 0.3125 |
| youngLD vs oldLD | **<0.0001** |
| youngHD vs youngLD | **<0.0001** |
| oldHD vs oldLD | **<0.0001** |
| oldHD vs youngLD | **<0.0001** |
| youngHD vs oldLD | **<0.0001** |

**Table S2**: Effect of parental age and density on fecundity in wheat.

Linear model: *fecundity ~ parental age * parental density*. Significant values (p < 0.05) are highlighted in bold.

|  | df | SS | MS | F | p |
| --- | --- | --- | --- | --- | --- |
| Effect of parental context | | | | | |
| Context | 3 | 15257 | 5086 | 80.15 | **<2.2e-16** |
| Effect of individual components of parental context | | | | | |
| Age | 1 | 977.78 | 977.78 | 15.41 | **1.20E-04** |
| Density | 1 | 11817.07 | 11817.07 | 186.24 | **3.65E-30** |
| Age x density | 1 | 2462.59 | 2462.59 | 38.819 | **2.84E-09** |
| Residuals | 194 | 12309.66 | 63.45 |  |  |

**Table S3:** Effect of parental context on egg survival and development rate. Results given are for a generalised linear model with binomially distributed errors

LR: likelihood ratio, significant values (p < 0.05) are highlighted in bold.

|  | | Wheat | | | FM | | |
| --- | --- | --- | --- | --- | --- | --- | --- |
|  |  | LR Chisq | df | p | LR Chisq | df | p |
| Effect of parental context | | | | | | | |
| Egg survival | Context | 23.93 | 3 | 2.58e-05 | 14.26 | 3 | **2e-03** |
| Egg development | Context | 39.02 | 3 | 1.71e-08 | 39.02 | 3 | **1.71e-08** |
| Effect of individual components of parental context | | | | | | | |
| Egg survival | Age | 21.78 | 1 | **3.06e-06** | 12.31 | 1 | **4.4e-04** |
|  | Density | 2.52 | 1 | 0.11 | 0.35 | 1 | 0.55 |
|  | Age x density | 9.67 | 1 | **1.8e-03** | 1.67 | 1 | 0.2 |
| Egg development | Age | 9.65 | 1 | **1.8e-03** | 9.65 | 1 | **1.9e-03** |
|  | Density | 7.07 | 1 | **7.9e-03** | 7.07 | 1 | **7.8e-03** |
|  | Age x density | 0.06 | 1 | 0.80 | 0.06 | 1 | 0.80 |

**Table S4:** Pairwise comparisons across contexts in each resource for the models show in Table S3, using the Benjamini-Hochberg correction for multiple tests. Significant corrected values (p < 0.05) are highlighted in bold. FM: finger millet.

|  | Comparison | p-value | Corrected p-value |
| --- | --- | --- | --- |
| Survival | wheat: youngHD vs oldHD | 1 | 1 |
|  | wheat: youngLD vs oldLD | 4.19e -06 | **2.51e-05** |
|  | wheat: youngHD vs youngLD | 0.16 | 0.31 |
|  | wheat: oldHD vs oldLD | 2.29e -03 | **1.10e-02** |
|  | wheat: oldHD vs youngLD | 3.27e-03 | **1.30e-02** |
|  | wheat: youngHD vs oldLD | 0.16 | 0.31 |
|  | FM: youngHD vs oldHD | 0.28 | 0.57 |
|  | FM: youngLD vs oldLD | 8.30e-04 | **4.98e-03** |
|  | FM: youngHD vs youngLD | 0.61 | 0.61 |
|  | FM: oldHD vs oldLD | 1.80e-01 | 5.39e-01 |
|  | FM: oldHD vs youngLD | 0.06 | 0.26 |
|  | FM: youngHD vs oldLD | 1.30e-02 | 6.50e-02 |
| Development rate | wheat: youngHD vs oldHD | 0.1 | **2.91e-02** |
|  | wheat: youngLD vs oldLD | 2.08e-02 | **4.16e-02** |
|  | wheat: youngHD vs youngLD | 4.09e-03 | **1.64e-02** |
|  | wheat: oldHD vs oldLD | 1.10e-03 | **5.50e-03** |
|  | wheat: oldHD vs youngLD | 2.49e-09 | **1.49e-08** |
|  | wheat: youngHD vs oldLD | 5.64e-01 | 5.64e-01 |
|  | FM: youngHD vs oldHD | 1.35e-01 | 2.70e-01 |
|  | FM: youngLD vs oldLD | 1.03e-02 | **4.12e-02** |
|  | FM: youngHD vs youngLD | 5.70e-03 | **2.85e-02** |
|  | FM: oldHD vs oldLD | 4.67e-02 | 1.40e-01 |
|  | FM: oldHD vs youngLD | 3.27e-06 | **1.96e-05** |
|  | FM: youngHD vs oldLD | 7.71e-01 | 7.71e-01 |

**Table S5:** Effect of parental context on number of eggs and their size, in dissected ovaries. Statistics for analysis performed excluding 3 potential outliers is given in parentheses. . Significant corrected values (p < 0.05) are highlighted in bold. Linear models: *egg number ~ parental age * parental density* and *Average egg size ~ parental age * parental density*

|  |  | SS | MS | F | p |
| --- | --- | --- | --- | --- | --- |
| Effect of parental context | | | | | |
| Number of eggs | Context | 6.08 (6.59) | 2.02 (2.2) | 0.51 (0.97) | 0.68 (0.41) |
| Egg size | Context | 3e-03 (3e-03) | 1e-03 (1e-03) | 4.28 (5.22) | **7e-03**  (**2e-03**) |
| Effect of individual components of parental context | | | | | |
| Number of eggs | Age | 1.8 (0.72) | 1.8 (0.72) | 0.45 (0.32) | 0.50 (0.57) |
|  | Density | 3.11 (5.41) | 3.11 (5.41) | 0.78 (2.38) | 0.38 (0.12) |
|  | Age x density | 1.17 (0.45) | 1.17 (0.45) | 0.3 (0.2) | 0.59 (0.66) |
|  | Residuals | 446.43 (249.88) | 3.98 (2.28) |  |  |
| Egg size | Age | 1e-03 (1e-03) | 1e-03 (1e-03) | 5.84 (7.169) | **0.02** (**8e-03**) |
|  | Density | 1.6e-03 (1.9e-03) | 1.6e-03 (1.9e-03) | 6.86 (8.47) | **0.01** (**4e-03**) |
|  | Age x density | 3e-05 (6e-06) | 3e-05 (6e-06) | 0.15 (0.03) | 0.7 (0.87) |
|  | Residuals | 0.02 (0.02) | 2.3e-04 (2e-04) |  |  |

**Table S6:** Effect of parental context and number of eggs in the ovary on egg size (ANCOVA results). Statistics for analysis performed excluding outliers is given in parentheses. Significant corrected values (p < 0.05) are highlighted in bold.

ANCOVA: *average egg size ~ parental context + number of eggs + parental context : number of eggs*

|  | SS | df | F | p |
| --- | --- | --- | --- | --- |
| Number of eggs | 5.6e-06 (2.7e-04) | 1 (1) | 0.02 (1.26) | 0.88 (0.26) |
| Context | 2.9e-03 (2.2e-03) | 3 (3) | 4.21 (3.42) | **7.5e-03** (**0.02**) |
| Number of eggs X context | 1.5e-03 (5.4e-03) | 3 (3) | 2.2 (0.84) | 0.09 (0.47) |
| Residuals | 0.02 (0.02) | 99 (93) |  |  |

**Table S7:** Effect of parental context on clutch protein and lipid. Statistics for analysis performed excluding outliers is given in parentheses. Results are for linear models. Significant corrected values (p < 0.05) are highlighted in bold.

Linear model: *clutch protein ~ parental age * parental density*

*Clutch lipid ~ parental age * parental density*

|  |  | df | SS | MS | F | p |
| --- | --- | --- | --- | --- | --- | --- |
| Effect of parental context | | | | | | |
| Clutch protein | Context | 3 | 12655 | 4221.8 | 1.32 | 0.28 |
| Clutch lipid | Context | 3 | 49296 (6133.5) | 16432 (2044.51) | 4.3 (7.78) | **0.009** (**0.002**) |
| Effect of individual components of parental context | | | | | | |
| Clutch protein | Age | 1 | 8857.25 | 8857.25 | 2.76 | 0.103 |
|  | Density | 1 | 1331.97 | 1331.97 | 0.41 | 0.52 |
|  | Age x  density | 1 | 2476.05 | 2476.05 | 0.77 | 0.38 |
|  | Residuals | 44 | 140720.89 | 3198.20 |  |  |
| Clutch lipid | Age | 1 | 16221.75  (1663.66) | 16221.74  (1663.66) | 4.24  (4.70) | **0.045**  (**0.035**) |
|  | Density | 1 | 11944.53  (585.023) | 11944.53  (585.02) | 3.123  (1.65) | 0.084  (0.20) |
|  | Age x  density | 1 | 21129.81  (3884.83) | 21129.81  (3884.83) | 5.52  (10.97) | **0.023**  (**0.0019**) |
|  | Residuals | 44  (42) | 168253.41  (14864.50) | 3823.94 | (353.91) |  |

**Table S8:** Effect of parental context on per egg protein and lipid. Statistics for analysis performed excluding outliers is given in parentheses. Results are for linear models. Significant corrected values (p < 0.05) are highlighted in bold.

Linear model: *protein per egg ~ parental age * parental density*

*Lipid per egg ~ parental age * parental density*

|  |  | df | SS | MS | F | p |
| --- | --- | --- | --- | --- | --- | --- |
| Effect of parental context | | | | | | |
| Protein per egg | Context | 3 | 1.64 (2.08) | 0.55 (0.69) | 0.32 (0.77) | 0.81 (0.52) |
| *Lipid per egg* | Context | 3 | 146.9 | 48.96 | 2.7 | 0.06  (**4e-03**) |
| Effect of individual components of parental context | | | | | | |
| Protein per egg | Age | 1 | 1.21  (1.11) | 1.21  (1.11) | 0.70  (1.23) | 0.41  (0.27) |
|  | Density | 1 | 0.24  (0.22) | 0.24  (0.22) | 0.14  (0.25) | 0.71  (0.62) |
|  | Age x density | 1 | 0.19  (0.74) | 0.19  (0.74) | 0.11  (0.83) | 0.74  (0.37) |
|  | Residuals | 44 (42) | 75.73  (37.71) | 1.72  (0.9) |  |  |
| *Lipid per egg* | Age | 1 | 139.44  (22.70) | 139.44  (22.70) | 7.64  (11.76) | **8.2e-03**  (**1.4e-03**) |
|  | Density | 1 | 0.51  (6.76) | 0.51  (6.76) | 0.03  (3.5) | 0.87  (0.07) |
|  | Age x density | 1 | 6.91  (0.70) | 6.91  (0.70) | 0.38  (0.37) | 0.54  (0.55) |
|  | Residuals | 44 (40) | 802.80  (77.21) | 18.24  (1.93) |  |  |

**Table S9:** Effect of parental context and fecundity on protein and lipid per egg. Results below are for an ANCOVA model. Statistics for analysis performed excluding outliers is given in parentheses. Significant corrected values (p < 0.05) are highlighted in bold.

ANCOVA: *protein per egg ~ parental context + fecundity + parental context : fecundity*

*Lipid per egg ~ parental context + fecundity + parental context : fecundity*

|  |  | SS | df | F | p |
| --- | --- | --- | --- | --- | --- |
| Protein | Context | 1.21 (0.14) | 3(3) | 0.30 (0.063) | 0.82 (0.98) |
|  | Fecundity | 21.58 (8.69) | 1(1) | 16.27 (11.79) | **2.4e-04** (**1.4e-04**) |
|  | Context x fecundity | 1.09 (0.99) | 3(3) | 0.27 (0.45) | 0.84 (0.72) |
|  | Residuals | 53.06 (28.02) | 40(38) |  |  |
| Lipid | Context | 139.25(12.82) | 3(3) | 2.7 (6.8) | 0.06 (**9.4e-04**) |
|  | Fecundity | 96.85 (48.08) | 1(1) | 5.63 (76.56) | **0.02** (**1.9e-10**) |
|  | Context x fecundity | 17.94 (6.52) | 3(3) | 0.35 (3.46) | 0.79 (**0.03**) |
|  | Residuals | 688.02 (22.61) | 40(36) |  |  |

**Table S10:** Effect of adult context (youngHD and oldHD) on protein and lipid. Results are for linear models. Significant corrected values (p < 0.05) are highlighted in bold.

Linear model: *protein ~ parental context*

*Lipid ~ parental context*

|  |  | df | SS | MS | F | p |
| --- | --- | --- | --- | --- | --- | --- |
| Protein | Parental context | 1 | 0.02 | 0.02 | 0.24 | 0.63 |
|  | Residuals | 28 | 2.82 | 0.10 |  |  |
| Lipid | Parental context | 1 | 5.3e-04 | 5.3e-04 | 0.79 | 0.39 |
|  | Residuals | 18 | 0.01 | 6.8e-04 |  |  |
